# An orthogonal and pH-tunable sensor-selector for muconic acid biosynthesis in yeast

**DOI:** 10.1101/229922

**Authors:** Tim Snoek, David Romero-Suarez, Jie Zhang, Mette L. Skjoedt, Suresh Sudarsan, Michael K. Jensen, Jay D. Keasling

**Affiliations:** Novo Nordisk Foundation Center for Biosustainability, Technical University of Denmark, Kgs. Lyngby, Denmark; Joint BioEnergy Institute, Emeryville, CA, USA; Biological Systems and Engineering Division, Lawrence Berkeley National Laboratory, Berkeley, CA, USA; Department of Chemical and Biomolecular Engineering & Department of Bioengineering, University of California, Berkeley, CA, USA

**Author notes:** Author of correspondence: Michael K. Jensen.

**Keywords:** transcriptional activator, biosensor, sustainability, evolution, metabolic engineering, yeast

## Abstract

Microbes offer enormous potential for production of industrially relevant chemicals and therapeutics, yet the rapid identification of high-producing microbes from large genetic libraries is a major bottleneck in modern cell factory development. Here, we develop and apply a synthetic selection system in *Saccharomyces cerevisiae* that couples the concentration of muconic acid, a plastic precursor, to cell fitness by using the prokaryotic transcriptional regulator BenM driving an antibiotic resistance gene. We show the sensor-selector does not affect production, and find that tuning pH of the cultivation medium limits the rise of non-producing cheaters. We apply the sensor-selector to selectively enrich for best-producing variants out of a large library of muconic acid production strains, and identify an isolate that produced more than 2 g/L muconic acid in a bioreactor. We expect that this sensor-selector can aid the development of other synthetic selection systems based on allosteric transcription factors.

## Introduction

In order to realize a bio-based economy, metabolic engineering aims to develop microbes that can convert inexpensive, renewable feedstocks into valuable products.^1^ Initial genetically-engineered strains, however, regularly need to be further optimized before their performance meets industrial demands on titers, rates and yields. Currently, decreases in DNA synthesis costs and the expansion of genome engineering tools allow for cost-effective building of large libraries of cell factory designs.^2,3^ However, since the vast majority of chemicals targeted for overproduction in microbes are not coupled to easy selectable phenotypes, evaluation of individual strains often relies on low-throughput analytical methods, severely challenging the turn-around time of the design-build-test-learn cycle.^4^

In recent years, development within synthetic biology has enabled the design and application of allosterically regulated transcription factors as biosensors.^5,6^ Such one-component regulators are abundantly present in prokaryotes,^7^ and can convert intracellular concentrations of otherwise inconspicuous chemicals of interest into easily measurable outputs, such as fluorescence (sensor-reporters) and antibiotic resistance (sensor-selectors) (reviewed by Rogers and colleagues^4^). Even in the yeast *Saccharomyces cerevisiae*, a well-established biotechnology workhorse, there is a large demand on improving current strains and generating yeasts that incorporate novel biosynthesis routes.^8^ To this end, a range of transcription factor-based biosensors that can aid the screening of yeast cell factory variants have been described,^9^ including sensor-reporters for detection of xylose,^10^ malonyl-CoA,^11^ *cis*, cis-muconic acid (CCM) and naringenin.^12^ In contrast to FACS-based evaluation of genetic libraries using sensor-reporters, sensor-selectors can be used to screen or select large libraries in high-resolution by simple and inexpensive coupling of chemical abundance with a growth-selectable phenotype. In prokaryotes, sensor-selectors have been widely used to select best-performing microbial strains or to evolve microbes,^13–15^ but also in yeast there are a few examples of the coupling of production to expression of auxotrophic marker genes.^16,17^

Previously, we have shown that transcriptional activators belonging to the LysR-type transcriptional regulator (LTTR) family can successfully be transplanted into yeast and applied as small-molecule sensor-reporters.^12^ One of the sensor-reporters, BenM, enabled expression of GFP correlated to *in vivo* CCM production. CCM is a platform chemical that can be converted into adipic acid or terephtalic acid, which can be further polymerized into numerous plastics.^18^ Whereas the highest CCM titer to date has been ascribed to *Escherichia coli*,^19^ from a process point of view producing CCM in a low-pH tolerant organism such as *S. cerevisiae* is of great interest. Rational engineering^20–22^ as well as evolution^23^ have been applied to establish and improve CCM production in yeast. Notably, Leavitt and co-workers used a synthetic reporter promoter inducible by aromatic amino acids (AAAs) to drive the expression of an antibiotic resistance gene and evolve a strain with an increased pool of endogenous AAAs.^23^ Following two consecutive rounds of EMS mutagenesis and adaptive laboratory evolution for approx. 600 h, the authors identified a strain producing 2.1 g/L CCM.

In order to design and apply faster and more simple sensor-selector systems based on small-molecule binding transcriptional activators, we reengineered our previously identified CCM sensor-reporter design into a sensor-selector. First, we determined the optimal design for the sensor-selector, taking into account parameters such as biosensor expression level and dynamic range. Second, we showed that the sensor-selector does not affect performance of yeast engineered to produce CCM. Third, we demonstrated that tuning pH of the medium can be used to minimize the rise of fast-growing, yet low-producing, cheaters. Finally, we applied the sensor-selector to enrich for best-producing strains out of a large library of CCM production strains. We also showed that our library contained an isolate able to produce more than 2 g/L CCM, on par with highest reported titers, and with higher productivity. To our knowledge, this is the first report on an orthogonal synthetic selection system in yeast driven by antibiotic resistance, which allows for rapid identification of best-producing cell factory variants from large strain libraries.

## Results and Discussion

### Design and characterization of a CCM sensor-selector

Previously we carried out a multi-parametric analysis in order to develop a CCM biosensor based on the LysR-type transcriptional regulator BenM transplanted from *Acetinobacter* sp. ADP1 into *S. cerevisiae.*^2^ Here, we set out to develop a sensor-selector based in *S. cerevisiae* to couple chemical production to growth by replacing the reporter gene from our previous study with the *KanMX* gene, a widely used marker conferring resistance to the antibiotic G418.^24^

In order to identify an optimal sensor-selector design supporting CCM-dependent growth under selective conditions (i.e. G418), we first compared the growth rates of yeast strains harboring the selector combined with no BenM, BenM expressed from the *TEF1* promoter, BenM expressed from the *REV1* promoter, and a previously identified^12^ triple BenM mutant (BenM*) expressed from the *REV1* promoter. Each of the four strains was pre-cultured in medium with or without CCM, and then subcultured into medium with the same composition with or without G418. We found that all four strains grew well in medium without selection, irrespective of the CCM concentration (Fig.1A). Contrastingly, without BenM, no growth was observed under selective conditions, whereas high BenM expression led to constitutive growth, even in the presence of G418 and absence of CCM. Finally, while low BenM expression showed modest CCM-dependent growth under selective conditions, the strain expressing BenM* showed pronounced CCM-dependent growth in the presence of G418. Therefore, REV1p-BenM* driving expression of the selector was chosen as the optimal design. For this design, we found that in medium without CCM or <40 mg/L CCM, no growth was observed under selective conditions (G418 present), whereas when grown in 80 – 200 mg/L CCM, this strain showed CCM-dependent tuning of the growth rate (Fig.1B). Taken together, these data demonstrate that this sensor-selector design has the potential to couple production of CCM to host growth.

**Figure 1:**
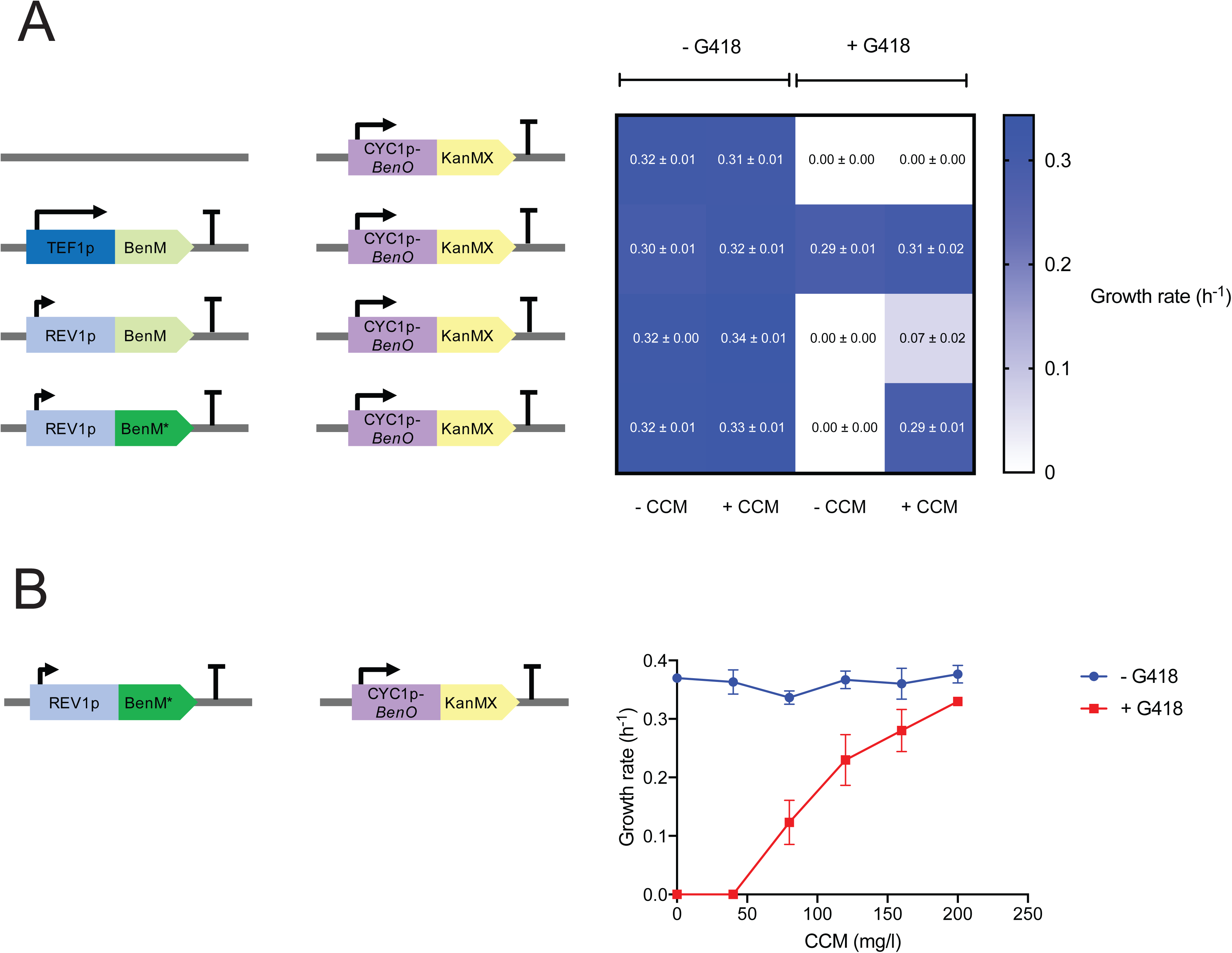
Characterization of biosensor designs for growth-coupled selection. (**A**) Four different strains harboring the selector gene (CYC1p_*BenO*-KanMX) and no, high (TEF1p), low (REV1p) expression of wild-type BenM, or low expression (REV1p) of a BenM triple mutant (BenM*). Cells expressing either of these four designs were pre-cultured in rich medium with or without 200 mg/L CCM, followed by subculturing into medium with the same composition with or without addition of 200 mg/L G418. Growth was monitored during 24 h. Means and standard deviations of growth rates based on biological triplicates are indicated in the heatmap. (**B**) The optimal sensor-selector design was tested in detail to determine the dose-response curve both in the presence and absence of G418. Growth rates are shown as mean ± s.d. from three (n = 3) biological replicates.

### Sensor-selector validation in production strain

Next, we aimed to investigate if the sensor-selector would be functional in yeast engineered to produce CCM from a 3-step heterologous biosynthetic pathway (Fig. 2A).^20,21^ We previously introduced this pathway into yeast and measured CCM production of individual variants differing in the number of integrated cassettes containing *Kp*AroY.B and *Kp*AroY.Ciso, genes encoding subunits of the rate-limiting enzyme AroY, which controls the conversion of protocatechuic acid (PCA) to catechol (see Fig. 2A and Methods).^12^ We chromosomally integrated the sensor-selector in one of those strains. We found no significant difference in growth rate between the original CCM-producing strain (CCM pathway) and the strain further engineered to express the sensor-selector (CCM pathway + sensor-selector) under non-selective conditions, though the CCM production strain grows significantly slower than wild-type CEN.PK (t-test, p<0.05), underscoring the growth burden of the production pathway (Fig. 2B). As expected, the production strain without the sensor-selector was not able to grow in selective medium, whereas the production strain with the sensor-selector showed robust growth in the presence of G418 (Fig. 2B). Moreover, there was no significant difference in CCM titer between the two strains (Fig. 2C). Taken together, these results show that the sensor-selector confers a growth-selectable phenotype when introduced to CCM-producing yeast without affecting CCM production.

**Figure 2:**
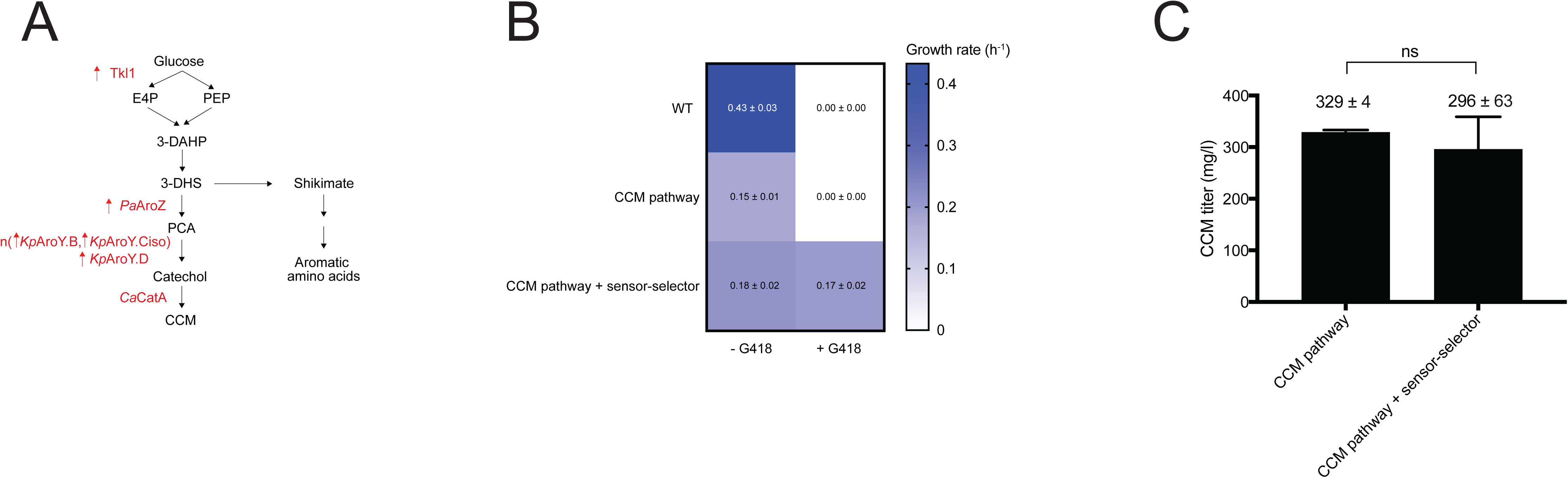
Biosensor-selector does not compromise cell factory performance. **(A)** A 3-step heterologous CCM pathway was built into yeast, comprising 21 single-copy expression of *PaAroZ*, *Kp*AroY.D, and *CaCatA* and multi-copy integration of a cassette expressing *Kp*AroY.B and *Kp*AroY.Ciso. Together with overexpression of *TKL1* this strain produces 300 mg/L CCM (Fig. 2C). (**B**) The sensor-selector was integrated into the CCM-producing strain (CCM pathway + sensor-selector) and its growth was compared to the baseline CCM-producing strain (CCM pathway) and wild-type CEN.PK. All three strains were cultured in medium with or without 200 mg/L G418, and growth monitored during 24 h. Mean and standard deviation of growth rates based on biological triplicates are indicated in the heatmap. (**C**) CCM titer was measured for the two CCM production strains after 72 h of cultivation. Mean and s.d. from four (n=4) biological replicates are shown. ns = not significant as evaluated by t-test.

### pH tuning of the sensor-selector system

One of the major considerations for bulk screenings of large diverse populations of cell factory variants is the rise of false-positives; i.e. cells that do not produce the compound of interest but are still able to thrive under selective conditions.^14,23^ This is especially relevant for biosynthetic pathways where production confers a growth burden, as observed for CCM (Fig. 2B). Due to the fact that protonated CCM can passively diffuse across the yeast cell membrane,^12^ we expected that one prominent way for cheaters to arise and be isolated from large genetic screens, would be for non-producing fast-growing cells to take up CCM secreted by slow-growing producing cells, resulting in sensor-selector activation. CCM is a weak acid with a pKa of 3.57, and for this reason we hypothesized that tuning the pH of the growth medium could control the rise of cheaters. In order to test this hypothesis, co-cultures of a CCM production strain (‘sender strain’) and a non-producing strain (‘receiver strain’) were performed. The receiver strain contains our previously described CCM sensor-reporter (REV1p-BenM* driving expression of yEGFP reporter gene),^12^ as well as a plasmid expressing RFP from the constitutive *TEF2* promoter (Fig. 3A). Co-cultures were performed in medium with pH 4.5 or pH 6 in three different sender:receiver starting ratios: 0:100, 90:10 and 99:1. After 24 h of co-culturing the fraction of the population consisting of the receiver strain (RFP), and the sensor-controlled reporter gene activity (GFP) in those cells was determined using flow cytometry. As inferred from percentages of RFP expressing cells, we found that population distributions looked similar for pH 4.5 and pH 6 (Fig. S1). However, for cells cultivated at pH 6, limited induction of the CCM-inducible sensor-reporter was observed, whereas for cells cultivated at pH 4.5, the biosensor activity was induced 3- to 8-fold approximately, depending on sender:receiver starting ratios (Fig. 3B and Fig. 3C). These data show that pH of the growth medium can control the degree of passive diffusion of CCM into non-producing cells, and that pH can provide a simple tuning parameter for bulk screening and selection of production strain libraries.

**Figure 3:**
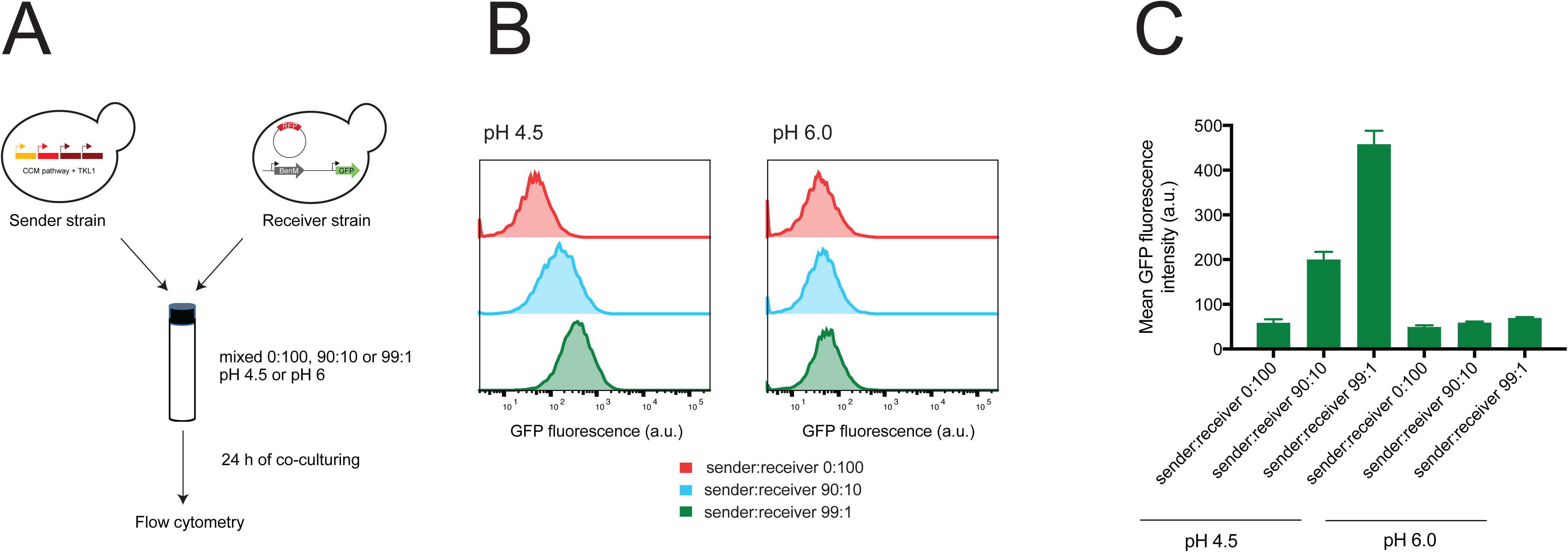
Safe-guarding selection from cheaters by pH control. (**A**) Outline of experimental set-up. A CCM-producing strain (sender) was co-cultured with a biosensor-reporter strain (receiver, no CCM production) in different inoculum ratios at pH 4.5 or pH 6. The receiver strain harbored a plasmid expressing RFP in order to identify biosensor cells by flow cytometry after 24 h of co-culturing. (**B**) After 24 h of culturing the mean GFP fluorescence intensity was measured in 10 000 RFP^+^-cells per co-culture. Histograms of representative populations are shown. (**C**) Mean GFP fluorescence intensity of RFP+-population is shown as mean ± s.d. for three biological replicates (n=3) per co-culture.

### High-throughput screening of a CCM production strain library

In order to determine whether the sensor-selector is able to enrich for high CCM-producing variants when grown in batch, we created a library of CCM-producing strains using a semi-randomized approach. As a starting strain, we used a strain overexpressing the *TKL1* gene encoding transketolase 1 in addition to the CCM biosynthetic pathway consisting of *Pa*AroZ, *Kp*AroY.D and *CaCatA*.^12^ This strain does not produce detectable amounts of CCM.^12^ We first integrated the sensor-selector into this strain, and following transformation of an expression cassette harboring *Kp*AroY.B and *Kp*AroY.Ciso for multi-copy integration into Ty4 sites,^25^ we obtained a library of approx. 10^4^ transformants (see Methods). Next, these transformants were pre-cultured in bulk in at pH 6, followed by inoculation of three different flasks with selective medium (i.e. 200 mg/L G418), as well as three flasks containing non-selective medium (Fig. 4A). Whereas the control cultures grew to saturation within approx. 30 h, the cultures growing under selective conditions needed more than 48 h to reach a similar cell density (Fig. 4B). Most importantly, whereas no CCM could be detected in the control cultures for any time point, a steady increase in CCM titer in the selective cultures, up to 275 ± 12 mg/L after 96 h, was observed (Fig. 4C), proving the power of the sensor-selector to robustly, and in high-throughput, enrich for CCM-producing variants.

**Figure 4:**
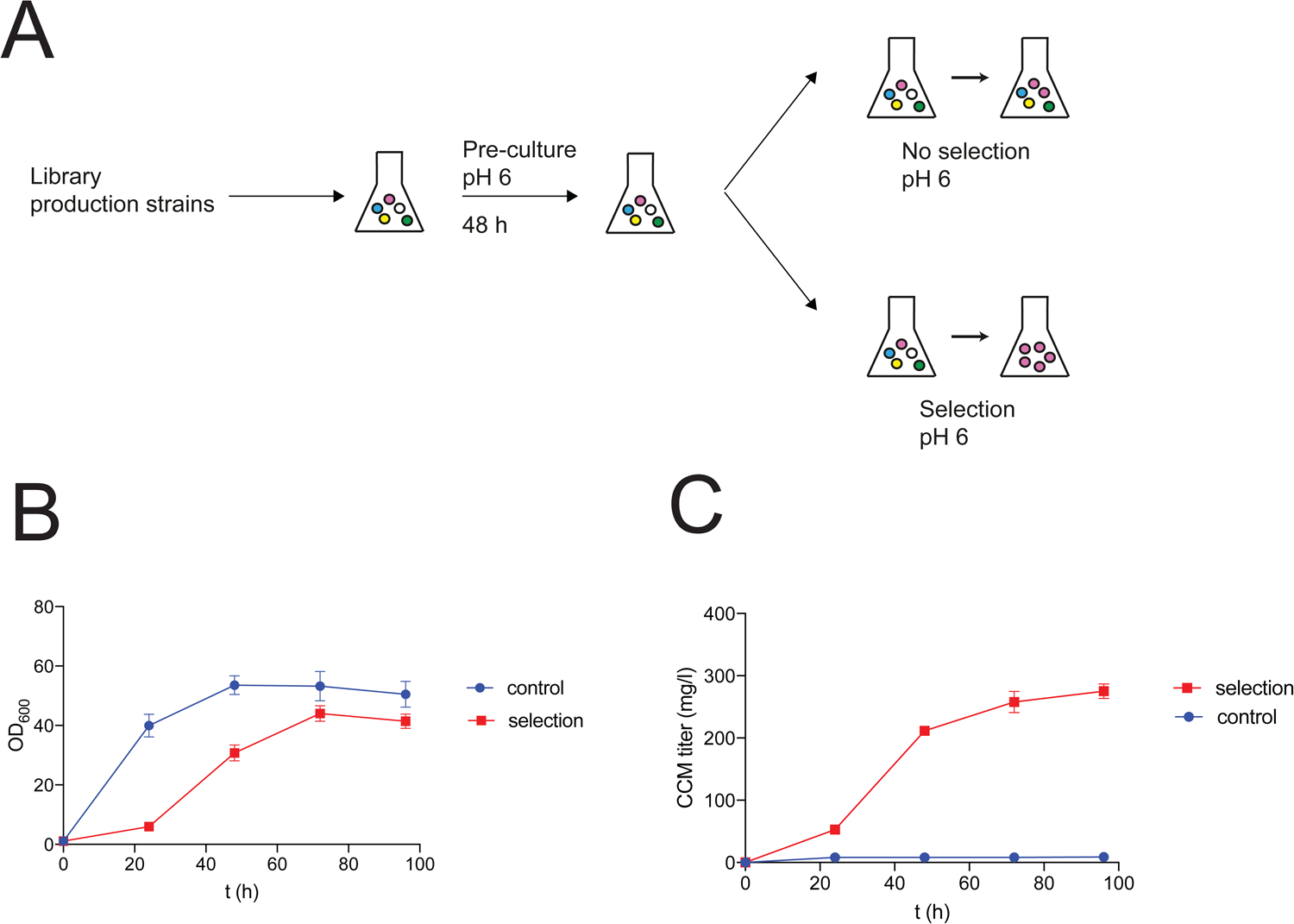
Pathway evolution using synthetic selection. (**A**) Experimental outline of multi-copy AroY library screening using the sensor-selector. The library was based on transformation of a construct consisting of *Kp*AroY.B and *Kp*AroY.Ciso targeting Ty4 sites into a strain containing the 3-step CCM production pathway and the sensor-selector. Library screening consisted of pre-culturing, followed by subculturing into selective medium as well as control cultures not containing G418. (**B**) OD600 values of strain library derived from 96 h cultivations under both selective and non-selective conditions. (**C**) CCM titers from library cultivations grown under both selective and non-selective conditions. For both (A) and (B), means and standard deviations for three biological replicates per cultivation condition are shown.

We suspected that before subculturing the cells into selective media, a proportion of the population may already consist of non-producing, fast-growing cells that have low copy numbers of *Kp*AroY.B and *Kp*AroY.Ciso. In order to verify this hypothesis, we characterized the starting population by measuring the growth with and without G418 of 89 single colonies isolated at the end of the pre-culture. From this, we observed a wide variation in growth rates in medium without G418, yet only two isolates were able to grow in the presence of G418 (Fig. S2). We measured the CCM production for the two G418-positive clones, as as well as for five isolates that were not able to grow in the presence of G418 spanning different growth rates (Fig. S2). Only the two G418-positive clones showed CCM production, whereas the remaining isolates did not produce detectable amounts of CCM (Table 1). These data show that right before applying selection, the library indeed consisted of a high proportion of fast-growing non-producing cells.

**Table 1:** Overview of growth rates (see Fig. S2), phenotype in G418-containing medium and CCM production in seven library isolates.

| isolate | phenotype | growth rate mineral medium (h<sup>-1</sup>) | CCM titer (mg/L) |
| --- | --- | --- | --- |
| 1 | G418- | 0.48 | 0.3 +/- 0.1 |
| 2 | G418- | 0.40 | 0.3 +/- 0 |
| 3 | G418- | 0.32 | 0.3 +/- 0 |
| 4 | G418- | 0.23 | 0 +/- 0 |
| 5 | G418- | 0.21 | 0 +/- 0 |
| 6 | G418+ | 0.15 | 379 +/- 3 |
| 7 | G418+ | 0.14 | 470 +/- 41 |

We next scaled-up CCM production of the two G418-positive isolates in bioreactors. In order to reach high titers, additional concentrated medium was spiked as soon as the CO_2_ production dropped, indicative of glucose depletion (see Fig. S3). Both strains reached high titers, with isolate 6 reaching 1905 ± 17 mg/L CCM (Fig. 5A), and isolate 7 producing 2028 ± 45 mg/L CCM (Fig. 5B). While not completely identical set-ups, for isolate 7, the productivity is almost doubled, while its titer is similar to the currently best-performing CCM production strain reported in literature.^23^

**Figure 5:**
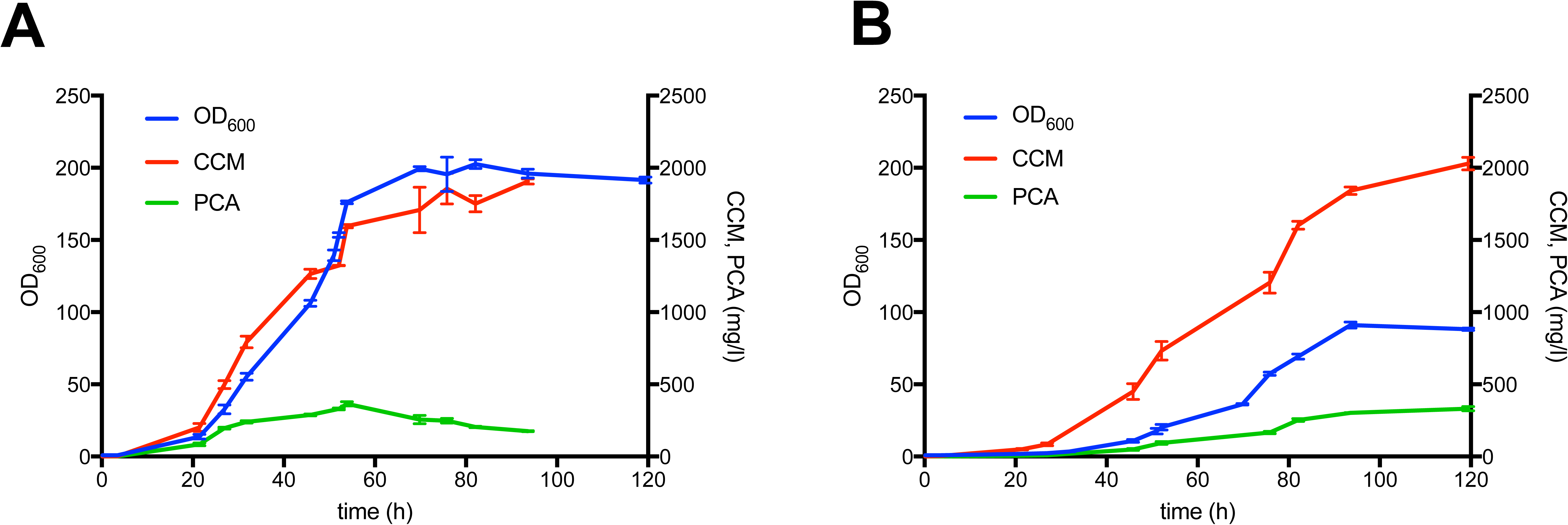
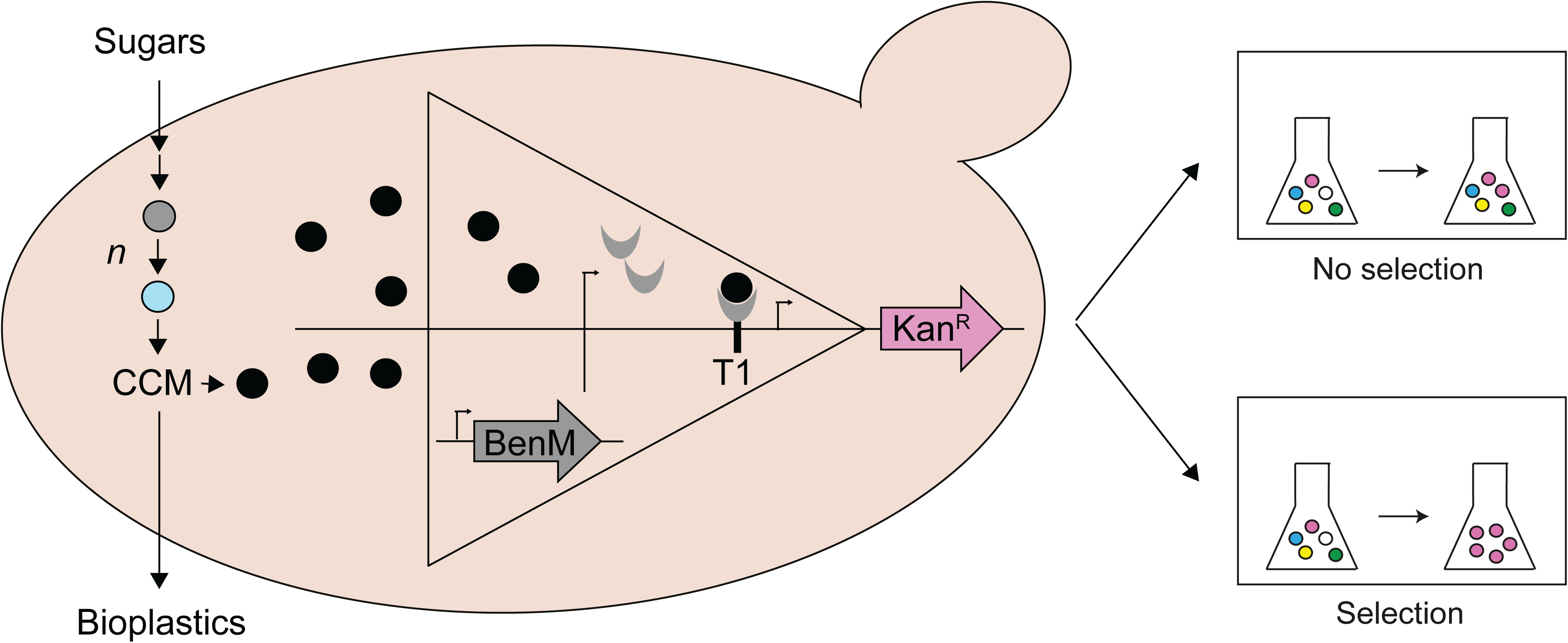
Bioreactor fermentations of selected CCM-producing strains. Biomass units (OD_600_), and titers of CCM and PCA from a repeated batch fermentation of isolate 6 (**A**) and isolate 7 (**B**) during a 120 h cultivation. For both (A) and (B), all values represent means and standard deviations from two biological replicates. See Figure S3 and Methods for details.

In summary, in this study, we designed, characterized and applied a fast and simple sensor-selector system in *S. cerevisiae* that directly couples the concentration of a chemical produced by a single cell to its fitness. Since BenM is part of the LTTR superfamily of small molecule-inducible prokaryotic transcriptional regulators,^26^ we envision that the sensor-selector system developed in this study could serve as a blueprint to develop high-throughput synthetic selection systems for a multitude of compounds regulated by LTTR-based transcription factors. Ultimately, this will significantly increase the turnaround time of the design-build-test-learn cycle for engineering future microbial cell factories.

## Methods

### Strains, chemicals and media

Yeast strains were grown on YPD, Synthetic Complete (SC) or mineral medium with urea (MMU). MMU was prepared as described previously^27^ with the exception that 2.3 g/L urea (Sigma, U1250) was used as a nitrogen source instead of ammonium sulphate, in order for G418 to be effective. Also, final pH was brought to 6.0, unless otherwise indicated. To test the response of non-producing cells to CCM (see further), *cis*, cis-muconic acid (Sigma, 15992) was always freshly dissolved in YPD medium, after which the pH of the medium was brought to 4.5 and filter-sterilized. CCM production strains were grown with selection for the destabilized uracil marker. *Saccharomyces cerevisiae* CEN.PK113-5A (MATa, *trp1 his3Δ1 leu2-3/112 MAL2-8^c^ SUC2*) and CEN.PK113-7D (wild type, *MATa MAL2-8^c^ SUC2*) strains were obtained from Peter Kötter (Johann Wolfgang Goethe-University Frankfurt, Germany). CCM production strain TISNO-11 was obtained from an EasyCloneMulti integration of *Kp*AroY.B and *Kp*AroY.Ciso as carried out previously.^12^ *Escherichia coli* strain DH5α was used as a host for cloning and plasmid propagation, and was grown at 37°C in Luria-Bertani medium supplemented with 100 μg/mL ampicillin. Phusion^®^ High-Fidelity DNA Polymerase or Phusion U Hot Start DNA Polymerase was used for PCR amplification according to manufacturer’s instructions.

### Plasmids and strain construction

An overview of plasmids used and constructed in this study is supplied in Supplementary Table 1. The lithium acetate method was used to transform yeast cells,^28^ followed by selection of transformants on synthetic drop-out medium (Sigma-Aldrich). For selection of strains transiently expressing KanMX and NatMX markers, 200 μg/mL G418 sulphate (Sigma, G8168) and 100 μg/mL nourseothricin dihydrogen sulfate (WERNER BioAgents, product no. 5.0), respectively, were added to the medium. Genomic integrations were achieved using EasyClone plasmids^27^ or marker-free EasyClone plasmids in combination with plasmids containing dominant markers on Cas9 and gRNA plasmids.^29,30^ Transformants were genotyped using oligonucleotides described in Supplementary Table 2. The resulting strains are listed in Supplementary Table 3.

The sensor and selector constructs (NotI digested pTS-5 and pTS-7) were integrated into EasyClone sites X-3 and XII-4, respectively, into strain ST2377 and TISNO-11 using pCfB2312 for Cas9 and pTS-9 for gRNA expression, which were subsequently cured off, generating strains DRS16 and TISNO-33, respectively. DRS16 formed the basal strain for the CCM production strain library. ST2377 was described previously^12^ and contains the dehydroshikimate DHS dehydratase from *Podospora anserina* (*P*aAroZ), the PCA decarboxylase genes from *Klebsiellapneumoniae* (*Kp*AroY.D), and the catechol 1, 2 dioxygenase CDO from *Candida albicans* (*CaCatA*). As carried out previously,^12^ we inserted multiple copies of a cassette containing *Kp*AroY.B and *Kp*AroY.Ciso using the EasyCloneMulti system^25^ into DRS16, and the library was obtained by adding all the cells post-transformation to a final volume of 25 ml SC-Ura, growing overnight, and storing cells in aliquots at -80 °C. Immediately after transformation, a defined number of cells was plated onto SC-Ura in order to determine the number of transformants as a proxy for library size.

### Library enrichment

Ten OD_600_ units of the library (>6600 coverage) were added to a total volume of 25 ml MMU pH 6 (starting OD_600_= 0.4) in 250 ml-Erlenmeyer flasks and grown for 48 h at 250 rpm, 30°C (pre-culture). After 48 h the pre-culture had reached OD_600_=7.5. At this stage a small portion of the pre-culture was plated for single colonies, of which 89 random clones were picked for growth rate determination. Biomass was harvested, centrifuged (5 min, 3000 g) and supernatant removed, and used to inoculate three selective cultures (25 ml MMU pH 6 + 200 mg/L G418) as well as three control cultures (25 ml MMU pH 6) to an initial OD_600_=1.0 and incubated at 30 °C, 250 rpm. The OD_600_ of the six cultures was determined on a daily basis for up to 96 h and every day 1 ml of broth was centrifuged and the supernatant saved for CCM quantification by HPLC.

### Growth rate determination

In different experiments the growth rate of yeast strains was determined. In order to assess the response curve to externally applied CCM of different sensor-selector designs, strains were grown overnight in 150 μl YPD per well (pre-culture). The next day, pre-cultures were subcultured 1:150 into either control medium (YPD pH 4.5) or YPD supplemented with CCM (40, 80, 120, 160 or 200 mg/L) at pH 4.5, followed by overnight growth. The next day, saturated cultures were diluted 1:150 into fresh medium with the same composition, with or without addition of 200 mg/L G418. CCM production strains were pre-cultured in SC-Ura medium overnight, and subcultured 1:150 into MMU with or without addition of 200 mg/L G418. Plates were sealed with Breathe-Easy^®^ sealing membrane (Sigma Z380059) and incubated at 30 °C in a platereader (BioTek ELx 808) with continuous shaking and OD630 measurements every 20 min for 24 h or 72 h. Growth rates were calculated using GATHODE software.^31^ For each strain and condition at least three biological replicates were measured.

### CCM production assays

For deep-well fermentations, strains were grown overnight in SC-Ura in a microtiter plate. The next day, the OD600 of the pre-cultures were measured, and strains were subcultured to starting OD_600_=1.0 (approximately 10^7^ cells/ ml) in 500 μl MMU in deep-well 96-well plates. After 72 h of incubation at 30°C, 300 rpm, the final OD_600_ was measured, cells were centrifuged (5 min, 3000 g), and the supernatant was used for HPLC quantification of CCM as described previously.^12^

For bioreactor cultivations, a procedure similar as previously described was followed.^23^ Two isolates, TISNO-219 (isolate 6) and TISNO-221 (isolate 7), were pre-cultured in 50 ml SC-Ura in duplicates in 250 ml-Erlenmeyer flasks overnight. The next day, the OD_600_ of each pre-culture was measured, and a portion of biomass was harvested in order to inoculate 1-L bioreactors (Sartorius, Gottingen, Germany) to starting OD_600_ of 1.0. The starting medium of each bioreactor was 500 ml MMU pH 6 containing 4% (w/v) glucose. During the cultivation, the stirring speed was maintained at 500 rpm and the dissolved oxygen level was kept above 20% by cascaded mixing of pure oxygen to the air stream (air input flow rate of 0.5 standard liter per minute). The pH was controlled at 6.0 by addition of 7 M NaOH, and the temperature was maintained at 30°C. At regular intervals, samples were withdrawn for OD600 measurement, afterwards centrifuged (5 min, 3000 × g, 4°C) and the supernatant kept for HPLC analysis to determine CCM, PCA and glucose levels. During the fermentation, the off-gas CO_2_ production was monitored continuously (Thermo Scientific Prima BT MS). Sterile fresh medium (containing 50 or 100 ml of 10X MMU medium) was pulsed after observing a significant drop in CO_2_ levels. In total 350 ml 10X MMU was added. Fermentations were performed for a total period of 5 days.

### Flow cytometry analysis of co-cultures

To determine the degree of biosensor-reporter activation in non-producing cells, co-cultures of sender strain TISNO-11 and receiver strain TISNO-31 were set up. Three single colonies of each strain were grown overnight in 3 ml SC-Ura-His-Leu-Trp. The next day the OD_600_ was measured, and co-cultures were started in MMU pH 6 or MMU pH 4.5 with a starting OD_600_ = 0.2. For each medium, sender and receiver strain were mixed in 0:100, 90:10 and 99:1 ratios in triplicates. After approx. 24 h cultures were diluted in PBS and analyzed on a BD Biosciences Aria (Becton Dickinson) with a blue laser (488 nm) to detect yeGFP and a yellow green laser (561 nm) to detect mKate2. FCS files were incorporated into FlowJo, and per replicate the mean GFP intensity of 10 000 RFP+ cells was determined after gating for single-cell events.

## Associated content

Supporting information: Supplementary Figures and Supplementary Tables (PDF).

## Author contribution

TS, MKJ and JDK conceived this project. TS and DRS designed the experiments and carried out experimental work. SS helped designing and carrying out repeated batch fermentations. MLS and JZ provided strains and materials. TS, DRS, MJK and JDK analyzed and interpreted the data. TS and MKJ wrote the paper, all authors assisted in this process.

## Notes

JDK has a financial interest in Amyris, Lygos, Constructive Biology, and Demetrix.

## Acknowledgements

We are grateful to the Novo Nordisk Foundation for funding. In addition, we thank Alicia Viktoria Lis for consultancy regarding HPLC analysis and Christoffer Knudsen for assistance with the repeated batch fermentations.

